# Using all available evidence to solve kinship cases

**DOI:** 10.1101/2025.05.03.652046

**Authors:** Thore Egeland, Franco Marsico

## Abstract

Kinship cases, ranging from standard paternity tests to complex disaster victim identifications, are typically evaluated using likelihood ratios (LR) based on forensic genetic markers. However, in some contexts, genetic information alone is not enough to reach conclusive results. This is common when establishing distant familial connections using large DNA-databases, or even in simple cases such as determining which individual is the parent and which is the child in a relationship pair. Although forensic practitioners frequently incorporate additional evidence (SE), such as age, biological sex, or phenotypic traits, in these cases, this integration typically occurs informally, without rigorous probability estimation, compromising procedural transparency and reliability. Here, we present a comprehensive methodological framework that formally synthesizes forensic DNA evidence (FDE) with SE through Markov chain models and customized transition matrices designed for various biological traits. This approach generates combined likelihood assessments expressed as LRs or posterior probabilities. Validation through simulated and real-world case studies demonstrates that systematic incorporation of SE improves resolution accuracy in kinship determinations. To facilitate adoption, we have implemented this methodology in `mispitools`, an open-source R package.

## 1 Introduction

This paper focuses on kinship analyses for forensic applications in missing person (MPI) and disaster victim identification cases (DVI). Kinship testing is generally addressed through genetic markers such as STRs and unlinked SNPs used as forensic DNA evidence (FDE). Although they are the standard for establishing biological relationships, forensic practitioners often face challenges in large-scale DNA database searches and mass grave scenarios, where FDE alone can lead to identification ambiguities [1, 2, 3, 4].

These ambiguities are particularly problematic in cases with insufficient DNA due to very distant relationships or degraded samples, leading to inconclusive results [2]. Also, even in straightforward parent-child testing, while FDE may strongly support a relationship, it cannot be used to respond which sample corresponds to the parent and which one to the child. Beyond being a simple question, it can become relevant when reconstructing genealogies or working with large mass graves [5].

Traditionally, inconclusive cases are addressed using supplementary evidence (SE), including age, biological sex, pigmentation, and physical traits [6, 7]. SE is commonly stored in large databases at international (e.g., Resolve platform and ICRC’s AM/PM database [8, 9]) and national levels (e.g., Colombia’s SIRDEC [10, 11], Argentina’s dictatorship database, among others [12]). However, SE is typically used informally without probability reporting [6, 12, 13, 14, 15], and no unified framework exists for computing likelihood ratios (LRs) across kinship problems, from parent-child relationships to large-scale MPI/DVI cases. The LR provides a transparent way to report the statistical weight of evidence by expressing the probability of observations under competing propositions and is recommended for forensic interpretation by ISO 21043 [16].

Figures 1 and 2 illustrate two pedagogical scenarios that serve as building blocks for our framework’s application to larger DNA database search problems. Figure 1 shows a parent-child relationship with unclear directionality, where *directional data* such as age differences can establish parental roles. This is a common scenario in burial or mass-grave [17] recovery and also in genealogical reconstructions using DNA databases [18]. The need to determine directionality also applies to grandparent-grandchild relationships, where small age differences between individuals can be used to identify false positives, commonly observed in database searches [2]. Figure 2 presents a DVI case [14] in which victims *V*_1_ and *V*_2_ must be matched to missing persons *M*_1_ and *M*_2_ based on their relationship with the reference *R*_1_. FDE alone cannot determine if *V*_1_ corresponds to *M*_1_ or *M*_2_, the same for *V*_2_. Therefore, *comparison data* (physical and pigmentation traits, among others) are commonly used to solve these cases, though direct matching [6].

**Figure 1.**
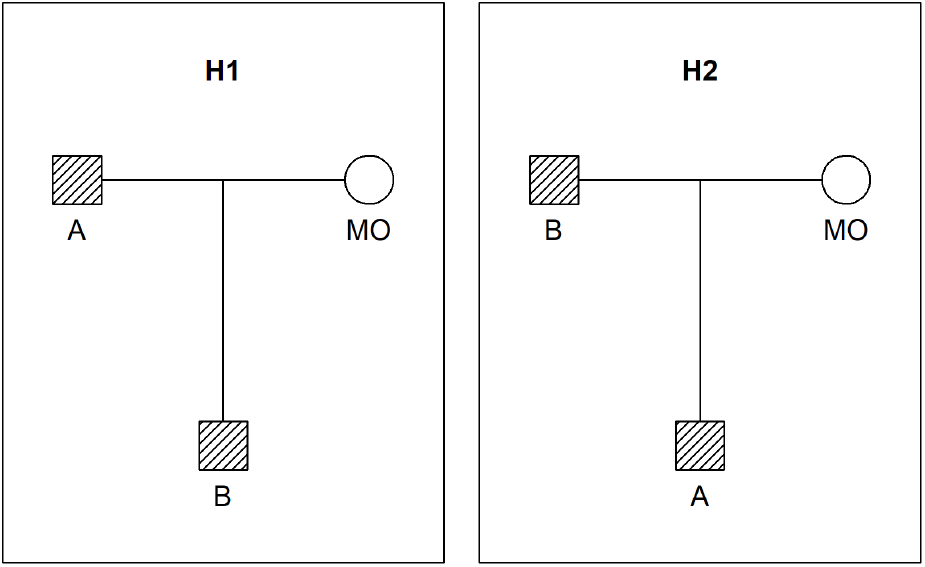
Pedigree scenario. Father - son hypotheses. Squares represent males and circles females. A and B are genotyped (dashed). *H*_1_ and *H*_2_ represent the two hypotheses which switch the position of A and B, therefore defining two pedigrees, 𝒫_1_ and 𝒫_2_.

**Figure 2.**
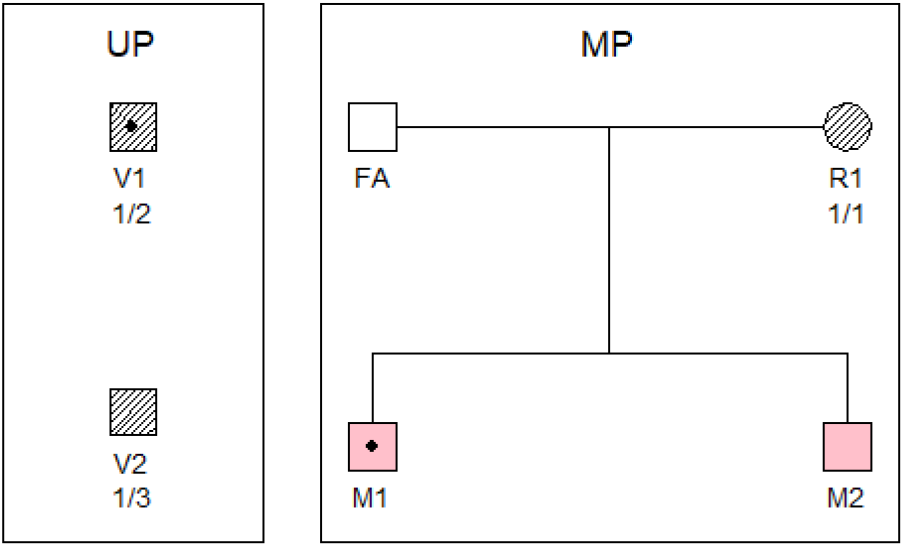
DVI case. Genotypes are available for the unidentified persons (samples) *V*_1_, *V*_2_ and the reference *R*_1_. *V*_1_ and *M*_1_ share a feature, indicated by a dot, not seen in *V*_2_ and *M*_2_.

Our primary objective is to provide a statistical framework that systematically combines FDE with directional and comparison SE data using proper probabilistic treatment. We formulate a unified model that computes LRs across scenarios from simple cases to complex missing person identifications through a DNA database search.

## 2 Data and methods

### 2.1 Hypotheses

Consider first pedigree cases; Figure 1 shows an example. Then, there are typically hypotheses *H*_1_ and *H*_2_ specifying that the individuals are related according to pedigrees 𝒫_1_ or 𝒫_2_, respectively. Generally, there could be more hypotheses, say *H*_3_ and *H*_4_, specifying that, for example, A is the uncle of B or vice versa. However, in DVI applications, there are generally many more hypotheses. For example, those corresponding to the case presented in Figure 2. The seven possible solutions to the identification problem are listed along with likelihoods and LRs, explained in the following sections. A hypothesis, referred to as an assignment *a*, for the DVI problem we are addressing, is a one-to-one correspondence between a subset of *𝒱* = *{V*_1_, …, *V*_*s*_*}* and a subset of *ℳ* = *{M*_1_, …, *M*_*m*_*}*, typically with the requirement that all identifications are sex consistent. For example, in Table 1, a consistent assignment is *{V*_1_ = *M*_2_, *V*_2_ = *M*_1_*}*. Alternatively, we may write this more compactly as a tuple (*M*_2_, *M*_1_).

**Table 1:** The seven possible solutions to the DVI problem in Figure 2. Each setting is represented by a different *a* value. The likelihoods based on SE (*L*_*SE*_) and FDE (*L*_*F DE*_) are followed by the *LR*, using the last assignment where no one are identified as reference. The calculations are explained in Example 3.3.

| a | $V_1$ | $V_2$ | $L_{SE}$ | $L_{FDE}$ | LR<sub>:.7</sub> |
| --- | --- | --- | --- | --- | --- |
| $a_1$ | $M_1$ | $M_2$ | $(1 - m) \cdot (1 - m)$ | $\frac{1}{2}p_2 \cdot p_3 \cdot p_1^2$ | $\frac{(1-m)^2}{\alpha\beta} \frac{1}{8p_1^2}$ |
| $a_2$ | $M_2$ | $M_1$ | $m \cdot m$ | $\frac{1}{2}p_2 \cdot p_3 \cdot p_1^2$ | $\frac{m^2}{\alpha\beta} \frac{1}{8p_1^2}$ |
| $a_3$ | $M_1$ | * | $(1 - m) \cdot \beta$ | $p_2 \cdot 2p_1p_3 \cdot p_1^2$ | $\frac{1-m}{\alpha} \frac{1}{2p_1}$ |
| $a_4$ | $M_2$ | * | $m \cdot \beta$ | $p_2 \cdot 2p_1p_3 \cdot p_1^2$ | $\frac{m}{\alpha} \frac{1}{2p_1}$ |
| $a_5$ | * | $M_1$ | $\alpha \cdot m$ | $2p_1p_2 \cdot p_3 \cdot p_1^2$ | $\frac{m}{\beta} \frac{1}{2p_1}$ |
| $a_6$ | * | $M_2$ | $\alpha \cdot (1 - m)$ | $2p_1p_2 \cdot p_3 \cdot p_1^2$ | $\frac{1-m}{\beta} \frac{1}{2p_1}$ |
| $a_7$ | * | * | $\alpha \cdot \beta$ | $2p_1p_2 \cdot 2p_1p_3 \cdot p_1^2$ | 1 |

### 2.2 Framework and notation

Here we introduce the general framework; specific models are detailed in the following subsections. The data in our framework consist of (*d, x, y*) where *d* are the genetic data, *x* the SE on the unidentified persons, and *y* the SE data on the missing persons. Given an assignment *a* (identifying a subset of the MPs as a subset of the UPs), the likelihood of *a* is *P* (*d, x, y*|*a*). This can be developed as:

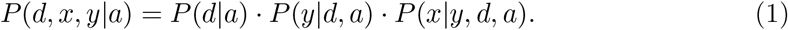

For the evaluation of *P* (*d*|*a*) see 2.4.1. Under the assumption that genetic data are independent of the SE data, we have *P* (*y*|*d, a*) = *P* (*y*). Considering *P* (*x*|*y, a*), it is evaluated as the product over *P* (*x*_*i*_|*y, a*). If *x*_*i*_ comes from an identified *M*_*j*_ according to *a* and there is corresponding UP data, we use the transition model (Equation (3) for one-dimensional data, or the binary error model in Section 2.4.3 for higher-dimensional data). Otherwise, we use priors with a Markovian assumption. The complete likelihood becomes:

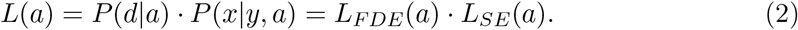

We assumed FDE and SE to be independent. This is valid for standard STRs or unlinked non-phenotype related SNPs. Denser SNP panels can introduce dependency within FDE and between FDE and SE [18] and are beyond the scope of the present work.

### 2.3 Priors

In some cases, we will present Bayesian approaches. This amounts to specifying prior probabilities *π*_1_, *π*_2_, … for the assignments, with, *π*_*i*_ *≥* 0, and ∑_*i*_ *π*_*i*_ = 1. In many cases, a flat prior is used, that is, *π*_1_ = *π*_2_ = · · ·. Parametric models are also available for pedigree priors [19, 20]. Finally, *empirical Bayes procedures* can be used to incorporate information [21]. An example of empirical Bayes is presented in 3.2. Priors are used for directional data in the examples.

### 2.4 Likelihood

#### 2.4.1 Forensic DNA Evidence

The evaluation of *P* (*d*|*a*) is performed using established methods detailed in [22, 23] and software packages such as `forrel` [24], which are part of the `pedsuite` libraries [25], or `Familias` [26].

#### 2.4.2 Supplementary Evidence

Consider a single feature *x*_*i*_ observed in an unidentified individual, *V*_*i*_, and its counterpart *y*_*j*_ in a missing person, *M*_*j*_. Features may be categorical, such as age and color, assuming multiple levels (*c >* 1), or continuous, such as age, height, among others. A floating bin approach [27] has been proposed to effectively handle continuous variables in forensic evaluations by conservatively assigning match windows that account for measurement uncertainty. Consequently, continuous variables can be recoded as binary match/nonmatch indicators. For example, assigning a match value of 1 when the age window of an unidentified person contains the age of a missing person, and a non-match value of 0 otherwise [13].

The conditional distribution of (*x*_*i*_ | *y*_*j*_, *V*_*i*_ = *M*_*j*_) is modeled by a matrix *c × c M* = (*m*_*st*_) where *m*_*st*_ *≥* 0 and 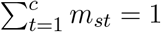. In other words, *M* is a transition matrix, similar to those used to model mutations in forensic genetics, as explained in Dawid et al [28]. In other words,

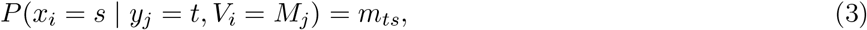

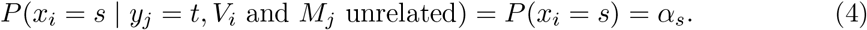

Note that we model conditionally on the data in the missing person. Thus, if there is uncertainty in the value in the missing person, this is not accommodated by the model (this topic is addressed in Section 4.1).

The transition matrix corresponding to Equation (3) may be written as

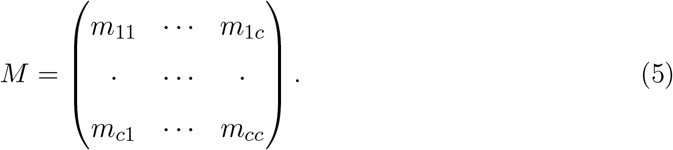

Knowledge of the feature may be reflected in the transition matrix. If we can assume that a possible error can only result in a classification to the closest value, we would use a band matrix like

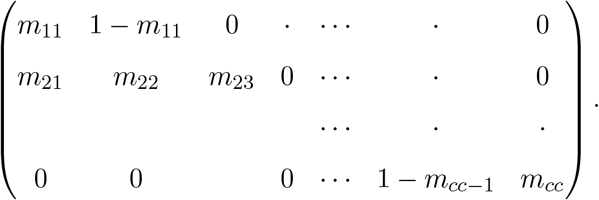

The likelihood based on the SE can be written

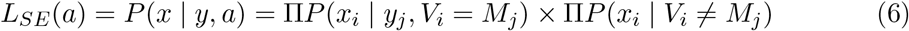

where the first product is over all pairs with *V*_*i*_ = *M*_*j*_, while the second product includes the remaining terms. The transition probabilities on the right-hand side of (6) are given by Equations (3) and (4). Note that (6) assumes independence between the feature values for UP-s. In the next section, we extend from one to several features.

#### 2.4.3 Modelling conditional dependency between traits

In practice, one feature will not suffice to resolve cases where some likelihoods based on FDE are identical. We therefore assume that *k* features have been observed for each UP and each M, and recorded, respectively, in *x*_*i*_ = (*x*_*i*1_, …, *x*_*ik*_) and *y*_*j*_ = (*y*_*j*1_, …, *y*_*jk*_). Consider the first product in Equation (6). We continue to assume that features between individuals are independent. Therefore, we need only consider *P* (*x*_*i*_ | *y*_*j*_, *V*_*i*_ = *M*_*j*_). We introduce variables to indicate if the feature values in UP and M coincide.

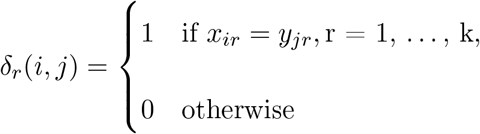

and *P* (*δ*_*r*_ = 0) = *ϵ*_*r*_. Note that we simplify to a binary error model rather than using more general matrices as in the previous section to limit the number of parameters. We assume that the features in an UP are independent given *V*_*i*_ = *M*_*j*_ and so

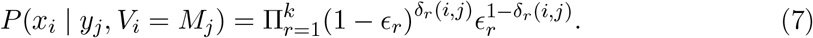

Note that *P* (*x*_*i*_ | *y*_*j*_, *V*_*i*_ = *M*_*j*_) = 1 if errors do not occur (we define 0^0^ = 1), i.e., when all epsilons are 0.

Consider next the likelihood under the hypothesis that *V*_*i*_ ≠*M*_*j*_. We assume that the feature vector *x*_*j*_ can be ordered to secure a Markovian dependence structure, i.e.,

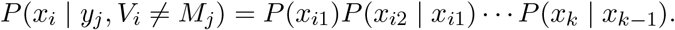

We estimate *P* (*x*_*is*_ = *u* | *x*_*i,s−*1_ = *v*) by (*N*_*u,v*_ + 1)*/*(*N*_*v*_ + *U*), where *N*_*u,v*_ is the observed frequency of the combination (*u, v*), *u* = 1, …, *U, v* = 1, …, *V*, and *N*_*v*_ is the total sample size when *x*_*i,s−*1_ = *v*.

### 2.5 Likelihood Ratios

The LR comparing the assignments *a* to *a*^*\**^ follows directly from Eq. (2):

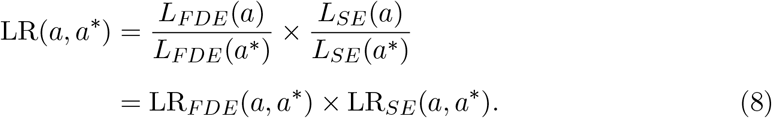

### 2.6 Posterior probabilities

There is nothing new in terms of converting LRs to posteriors:

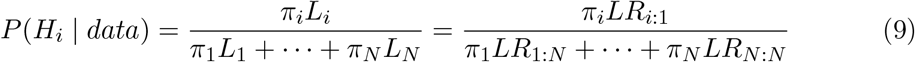

where *N* is the number of assignments and *LR*_*N*:*N*_ = 1.

### 2.7 Simulations and statistical power calculation

We define *TPR*(*T*) = *P* (*LR > T* |*H*_1_), *FPR*(*T*) = *P* (*LR > T* |*H*_2_), *TNR*(*T*) = *P* (*LR ≤ T* |*H*_2_), and *FNR*(*T*) = *P* (*LR ≤ T* |*H*_1_) to compute performance metrics. We use simulation-based evaluation [1, 2, 24] as described in Algorithm 1.

#### Algorithm 1

Evidence simulations

Buyer preferences for companies are influenced by factors extrinsic to the firm attributable to, and determined by, country-of-origin effects.

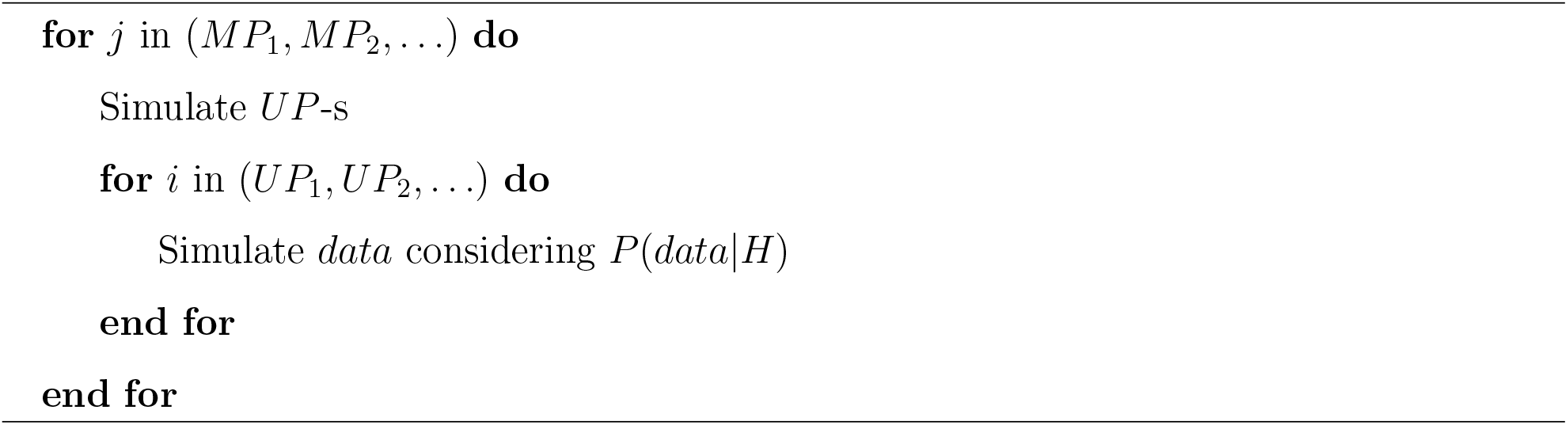

For each *M*_*j*_ in the database, the UP-s are simulated. Then, for each UP, *data* (that could be FDE or SE) is simulated considering *H*_1_ or *H*_2_ to be true. FDE simulations can be performed using the forrel package based on the methodology explained in [24], and SE simulation through mispitools package, explained in detail in [29].

### 2.8 The Balsac data

We use the BALSAC dataset [30], a genealogical repository from Quebec’s civil marriage records spanning four centuries. Data access is available upon request at https://balsac.uqac.ca. Further details are in Appendix B.

### 2.9 Implementation

We use R packages `forrel` [24], `pedsuite` [25] and `Familias` [19, 22] for FDE computations. `mispitools` [29] is used for SE models. Supplementary code (`usingAll.R`) is provided on https://github.com/MarsicoFL/mispitools/ for LR computations, simulations, and performance evaluation.

## 3 Results

In the first two examples, cases with directional data are presented. The comparison SE data are then introduced and exemplified in different settings. Finally, an example combining FDE and SE LRs is shown.

### 3.1 Directional data

In this subsection, we analyze some examples based on directional data, where SE can inform about possible kinship configurations.

#### Example 3.1.

We return to the example in Figure 1. This is one instance within a broad class of cases where the likelihood based on FDE remains unchanged under permutations of the genotyped individuals. In Figure 1, only two permutations are possible; with three full siblings, there are six.

The posteriors in (9) then reduce to the priors, *P* (*H*_*i*_ | data) = *π*_*i*_, since the likelihoods are identical. Distinguishing between hypotheses is only possible if prior information breaks the symmetry. As discussed in Section 2.3, priors can be obtained by: (a) direct specification, (b) a parametric model, or (c) empirical Bayes.

The first option is common in practice, e.g., if only one pedigree in Figure 1 is admissible. The parametric approach yields flat priors in symmetric settings and is not useful here. The empirical Bayes approach requires individual-level data and connects to the DVI framework.

Although resolving a single parent–child case may appear simple, in large-scale databases with multiple candidate matches, especially in multi-generational genealogies such as BALSAC [30], automating parentage recognition becomes valuable. Incorporating age can further help detect false positives. While rare for parent–child pairs, such false positives are more common in grandparent–grandchild cases [2], especially when using standard STR markers.

#### Example 3.2.

This case is exemplified in the BALSAC database context. Sample *A* is a highly degraded femur recovered from a clandestine grave; only fifteen mini-STR loci were successfully amplified. Sample *B* comes from a donor who stated he was *looking for a missing niece* from a half-brother but without knowing more pedigree details. Database matching shows that *A* and *B* are second-degree relatives consistent with the father of *A* and *B* being half-siblings, yet it is unclear whether the connection is through the father’s paternal line (*H*_1_) or maternal line (*H*_2_). Additional lineage markers cannot be obtained: the extract from *A* is exhausted, *A* is female (so Y-STRs are uninformative) and the quantity of DNA is insufficient for mtDNA re-amplification.

Civil registries such as BALSAC (Appendix B) show that roughly 65% of half-sibling pairs are paternal and 35% are maternal, giving the empirical prior

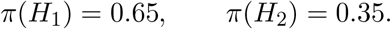

Because autosomal mini-STRs do not distinguish the two hypotheses,

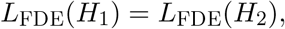

the posterior probabilities equal the prior values,

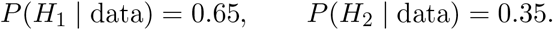

With laboratory options exhausted, this posterior becomes the sole quantitative guide for fieldwork: investigators first search among relatives in the father’s paternal branch and shift to the maternal branch only if that lead fails. It also helps to determine whether investing in additional analyses with dense SNP panels is warranted, assays that could provide further resolution but come at higher cost and may consume the limited DNA extract. Thus, even a coarse empirical-Bayes prior converts a genetically uninformative result into a targeted investigative strategy. This type of prioritization problem has previously been assessed through methodologies that only incorporate DNA data and is impractical here [24, 2, 31]. Therefore, SE becomes central to the prioritization.

### 3.2 Comparison data

In this subsection, we analyze a set of examples based on comparison data.

#### Example 3.3.

This example is based on Figure 2. We have a binary feature with value 1 (indicated by a dot in the figure), and value 2 (no dot) with frequencies *α* and *β* = 1 *− α*, respectively. The probability of misclassification is *m*, i.e.,

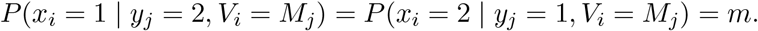

We see from the figure that the SE data is *x*_1_ = 1, *x*_2_ = 2, *y*_1_ = 1, *y*_2_ = 2 while the FDE is 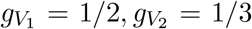 and 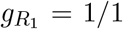. The possible assignments, the corresponding likelihoods and the LRs are given in Table 1. These calculations are based on the general likelihood given in (2) and the corresponding LR (8). We provide the details for the last line of the table. Regarding the SE, we find

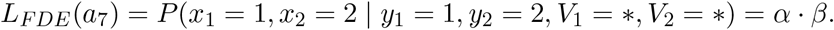

The conditioning on the missing persons is in this case irrelevant since they are unrelated to the unidentified persons. Moreover, we have assumed that the feature occurs independently in the UP-s, explaining the above equality. Turning to the FDE (STRs), we multiply the genotype probabilities of the genotyped individuals since they are unrelated in this case to arrive at

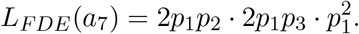

We can divide the assignments of Table 1 into three groups:

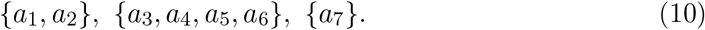

The assignments within the first two groups cannot be distinguished based on FDE alone.

We next give some numerical examples. Obviously, estimates of the parameters are then needed. Allele frequencies are obviously treated similarly as in standard forensic applications. Regarding, the feature frequency, *α* and *β* = 1 *− α*, reliable estimates may or may not be available. The hardest parameter to estimate is *m*, the probability of misclassification. If *m* = 0, corresponding to perfect feature classification, assignments *a*_2_, *a*_4_ and *a*_5_ are excluded. At the other extreme, *m* = 0.5, there is no information to distinguish within the three groups *{a*_1_, *a*_2_*}, {a*_3_, *a*_4_*}* and *{a*_5_, *a*_6_*}*. Table 2 provides a numerical example based on Table 1. In most practical cases, there will be enough forensic markers to distinguish *between* the three mentioned groups (10). The challenge is to distinguish within the groups and then we need more SE evidence as discussed in the next examples.

**Table 2:** LRs and posteriors calculated based on Table 1. Here *p* = (0.1, 0.2, 0.7), *α* = 0.8 and *m* = 0.1. The posterior is calculated as explained in Section 2.6.

| | $V_1$ | $V_2$ | combinedLR | LRSE | LRFDE | posterior |
| --- | --- | --- | --- | --- | --- | --- |
| a1 | M1 | M2 | 63.2812 | 5.0625 | 12.5000 | 0.6570 |
| a2 | M2 | M1 | 0.7813 | 0.0625 | 12.5000 | 0.0081 |
| a3 | M1 | * | 5.6250 | 1.1250 | 5.0000 | 0.0584 |
| a4 | M2 | * | 0.6250 | 0.1250 | 5.0000 | 0.0065 |
| a5 | * | M1 | 2.5000 | 0.5000 | 5.0000 | 0.0260 |
| a6 | * | M2 | 22.5000 | 4.5000 | 5.0000 | 0.2336 |
| a7 | * | * | 1.0000 | 1.0000 | 1.0000 | 0.0104 |

#### Example 3.4.

We illustrate how dependence between two phenotypic traits like hair and eye colour can affect the LR in a kinship context. Let **x** = (*x*_1_, *x*_2_) be the pigmentation traits, hair and eye colour respectively, of the unidentified person and **y** = (*y*_1_, *y*_2_) for the missing person. We apply the model presented in Section 2.4.3.

Joint frequency estimates for hair and eye colour can be obtained from large forensic DNA-phenotyping databases [7, 32]. Figure 3 depicts an example of the resulting probabilities for hair colour (1 = black, 2 = brown, 3 = red, 4 = blond) and eye colour (1 = brown, 2 = blue, 3 = green). Darker cells indicate more common trait combinations.

**Figure 3.**
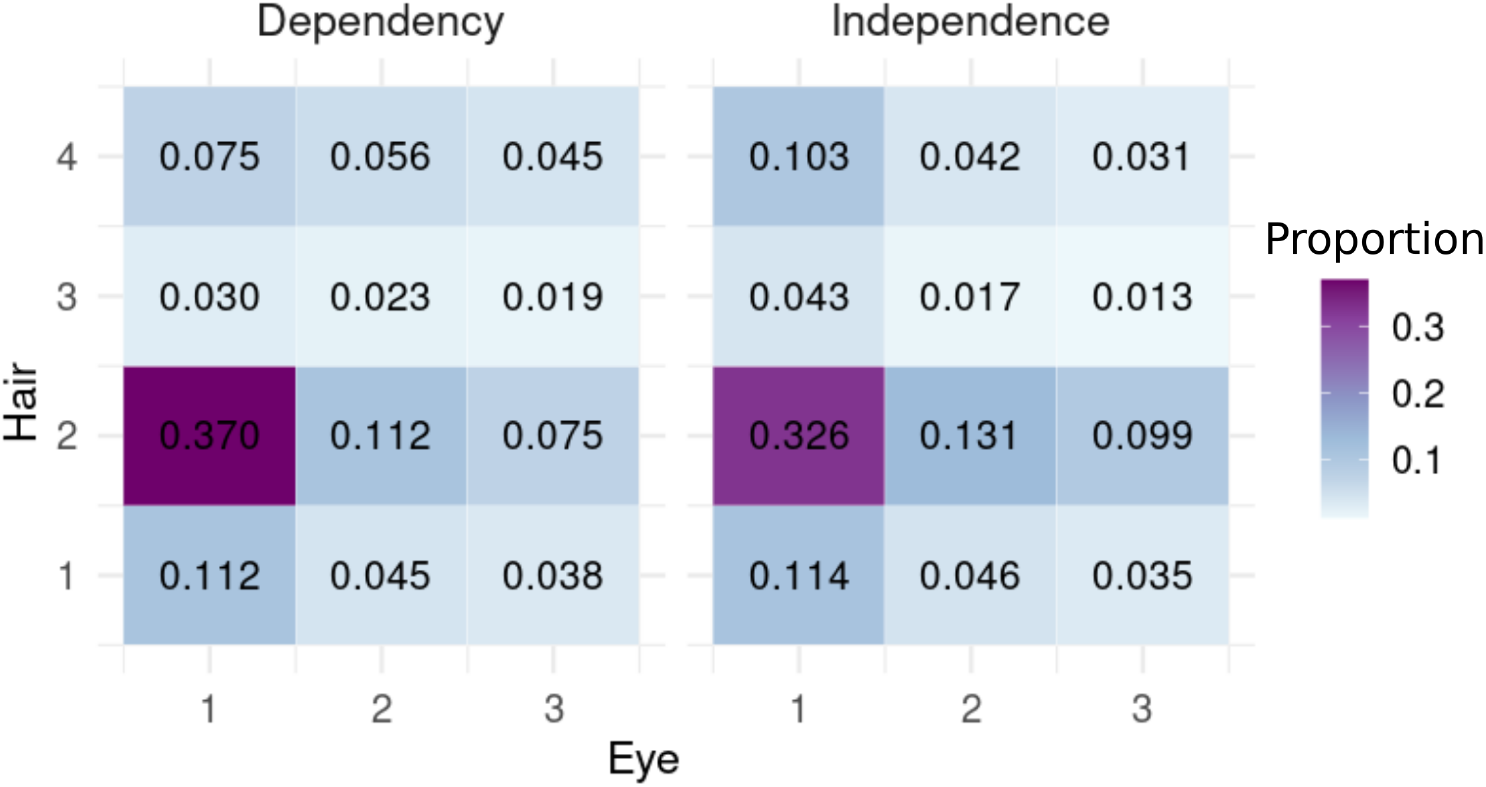
Joint frequencies 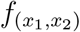 under *H*_2_, using (left) a conditional dependency model and (right) an independence model. Each cell corresponds to a combination (*x*_1_, *x*_2_), where *x*_1_ *∈ {* 1 = black, 2 = brown, 3 = red, 4 = blond *}* and *x*_2_ *∈ {* 1 = brown, 2 = blue, 3 = green *}*. The darkest cells represent the most frequent combination, here brown hair with brown eyes.

Under *H*_1_, each feature *x*_*i*_ is assumed to match its counterpart *y*_*i*_ with probability 1 *− ϵ*_*i*_ and to be misclassified with probability *ϵ*_*i*_. The LR comparing *H*_1_ to *H*_2_ is:

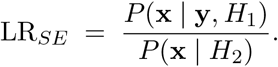

The denominator *P* (**x** | *H*_2_) follows either the dependency or independence assumption.

As an example, we consider a case where two most common pigmentation characteristics match between M and UP. For the dependency model,

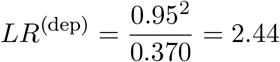

while independence gives

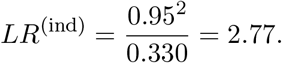

This means that not considering the dependency leads to an overestimation of the LR; the ratio independence/dependence is 2.77*/*2.44 = 1.14. In this case, overestimation is reasonable since ‘brown hair’ and ‘brown eyes’ are positively correlated. The denominator of the LR is then smallest for independence. Therefore, if we assume independence, we overestimate the power of observing this joint feature. In the extreme case of correlation 1, the second feature would contain no additional information if the first is observed. The features ‘brown eyes’ and ‘blond hair’ are negatively correlated and in this case the ratio is 0.075/0.103 = 0.73. In other words, ignoring dependence leads to underestimation. The effect of modeling dependence can be explored for different traits combinations, as shown in Table 3. It illustrates that for certain observed combinations, especially rare ones, the LRs can differ markedly between the two models.

**Table 3:**
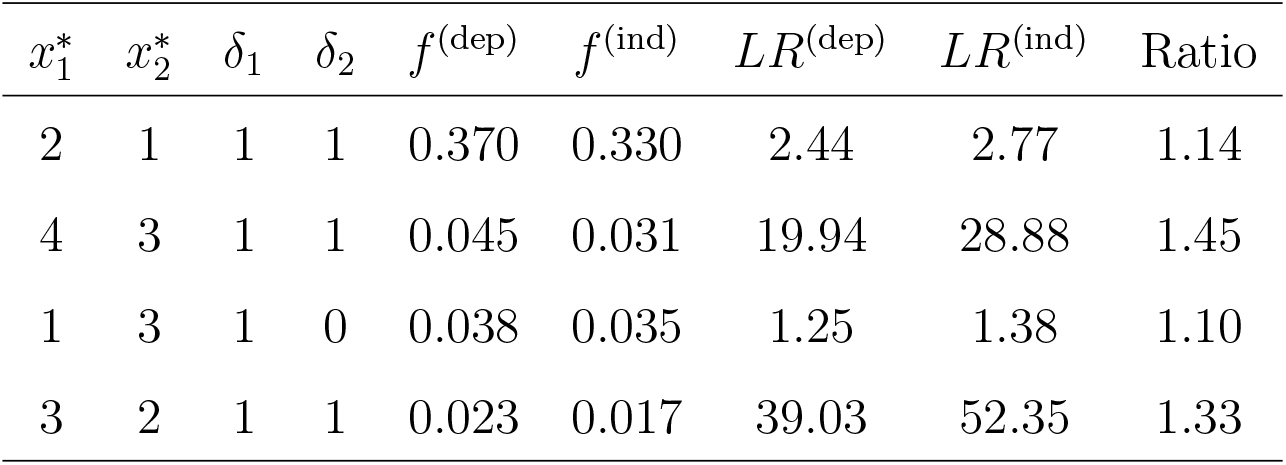
LR values under the dependency vs. independence models for four selected trait combinations. The columns *f* ^(dep)^ and *f* ^(ind)^ show the smoothed population frequencies under *H*_2_, with and without conditional dependence. The indicators *δ*_1_, *δ*_2_ specify which traits match under *H*_1_. The Ratio is obtained by dividing *LR*^(ind)^ by *LR*^(dep)^.

#### Example 3.5.

Power considerations have been useful for FDE [1, 2, 3, 4, 24, 31]. In this example, we extend the application to SE, facilitated by the provided LR models. Given the likelihood functions for *H*_1_ and *H*_2_, it is possible to evaluate *P* (*LR*_*SE*_ *≤ x*|*H*_1_) and *P* (*LR*_*SE*_ *≤ x*|*H*_2_). In particular, this approach allows researchers to:

i. determine the expected *LR* values under both *H*_1_ and *H*_2_ [33];
ii. assess the discrimination capacity of *LR* by analyzing the overlap between *P* (*LR*_*SE*_|*H*_1_) and *P* (*LR*_*SE*_|*H*_2_) [2];
iii. use simulations to predict which new evidence is most likely to enhance the method’s discrimination power [24, 31].

We analyse these properties using a missing person case from the Balsac database (see Appendix B). The subject (M) is a female, with an age at death of 40 ∓ 5 years) and brown hair. Population frequencies for age and sex are taken from the Balsac dataset, while hair color frequencies are based on published data [34]. A conservative error rate of *ϵ* = 0.05 is used for all models [6] and sensitivity analyses of all SE models are provided in Appendix C.

In total, 10,000 UPs were simulated under both *H*_1_ and *H*_2_. For example, under *H*_1_, approximately 95% of the simulated UPs were female (consistent with M), while under *H*_2_ the frequency was about 50%, matching the reference population. Similar simulations were carried out for hair color and age. For each simulated UP, an LR was computed using the described model and case, yielding LR distributions for each trait under both hypotheses. Also, combined LR distribution for all traits was obtained using the direct product between LRs.

Figure 4 illustrates these LR distributions. In Panel A, for instance, the blue bar (corresponding to *H*_1_) shows a higher frequency of cases with *Log*_10_(*LR*) = 0.28, whereas the brown bar (for *H*_2_) indicates that lower LR values are more common. Similar patterns are observed for hair color (Panel B), age (C) and combined (D).

**Figure 4.**
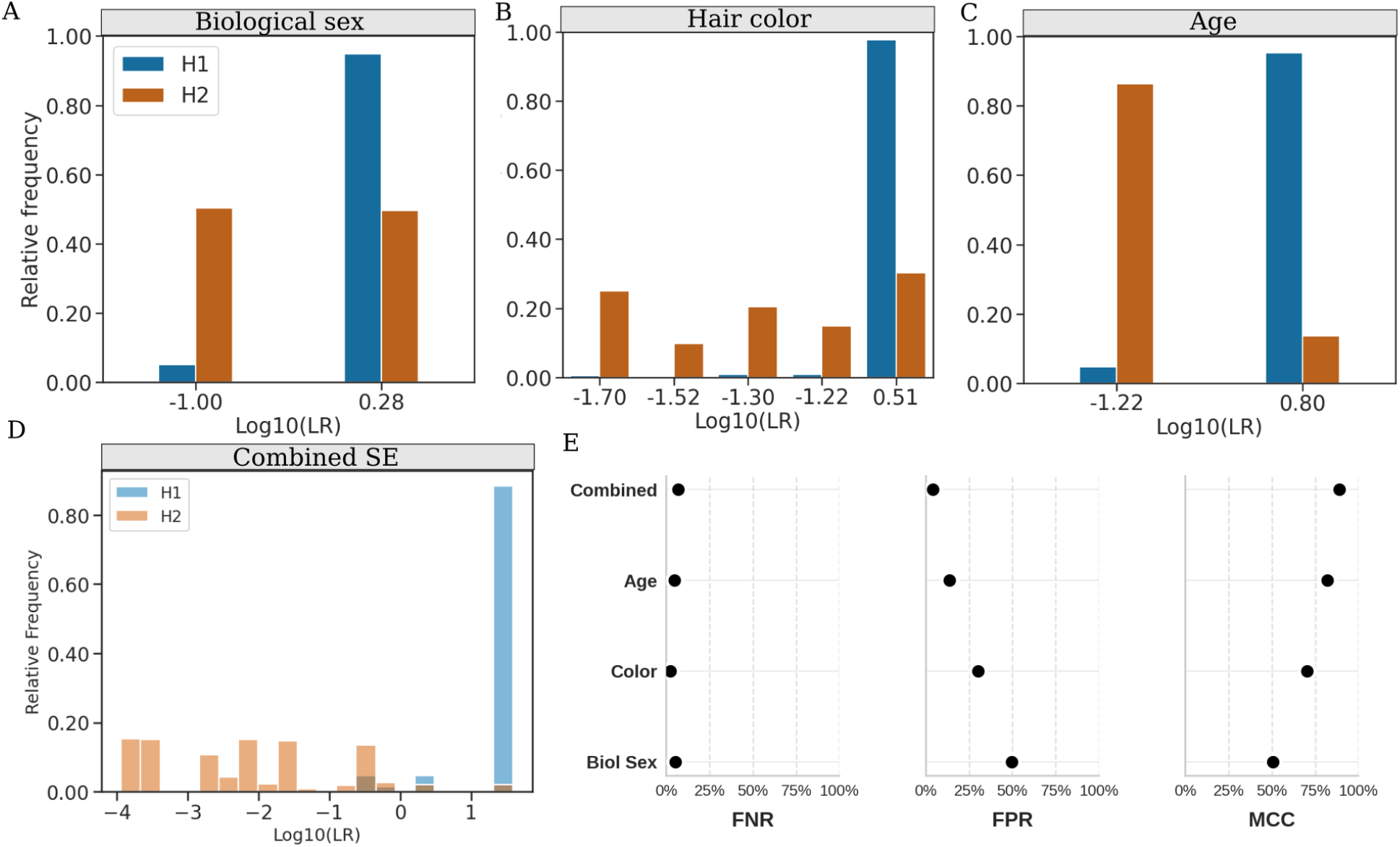
Panels A-C show the distributions of *Log*_10_ for three separate variables: biological sex, hair colour, and age, respectively, under the hypotheses *H*_1_ : UP is M (blue bars) and *H*_2_: UP is not M (brown bars). Panel D illustrates how combining these variables (under conditional independence) produces a sharper separation between LR distributions. Panel E shows performance metrics (FNR, FPR and Matthews Correlation Coefficient, MCC) compared for each approach, highlighting the improved discrimination achieved when multiple supplementary evidences are used. MCC delivers a single, balanced measure of overall accuracy that remains robust to class imbalance. LR threshold = 1 for all cases.

The results reveal a clear trend in the ability of SE variables to differentiate between *H*_1_ (UP is M) and *H*_2_ (UP is not M). As shown in panels A–C, Biological Sex provides the weakest discriminatory signal, with an MCC = 0.5049, meaning that nearly half of non-matching individuals could be incorrectly classified.

This aligns with expectations, given the limited number of sex categories and their relatively balanced distribution in human populations. Hair colour improves performance (FPR = 0.3018, MCC = 0.7027), likely due to its increased variability and predictive power in some populations.

However, Age stands out as the most informative single variable, with a notably low FPR (0.1361) and the highest MCC (0.8203) among individual traits.

The greatest improvement occurs when all SE variables are combined (Figure 4D). Although the FNR slightly increases (0.0692), the FPR drops dramatically to 0.0408, and the MCC reaches 0.8904, indicating a high classification performance.

### 3.3 Combining SE and FDE data

#### Example 3.6.

Finally, we investigated two pedigrees, each of which had low statistical power when relying solely on FDE (Figure 5). This implies that low LR values (*LR <* 1) can be obtained when *H*_1_ is true (*FNR*), and high values when *H*_2_ is true. The SE data are the same as those analyzed in the previous example. For FDE, 23 autosomal STR markers were considered. In Pedigree 1, the maternal grandmother and great-grandmother were available for genotyping; in Pedigree 2, only a paternal first cousin.

**Figure 5.**
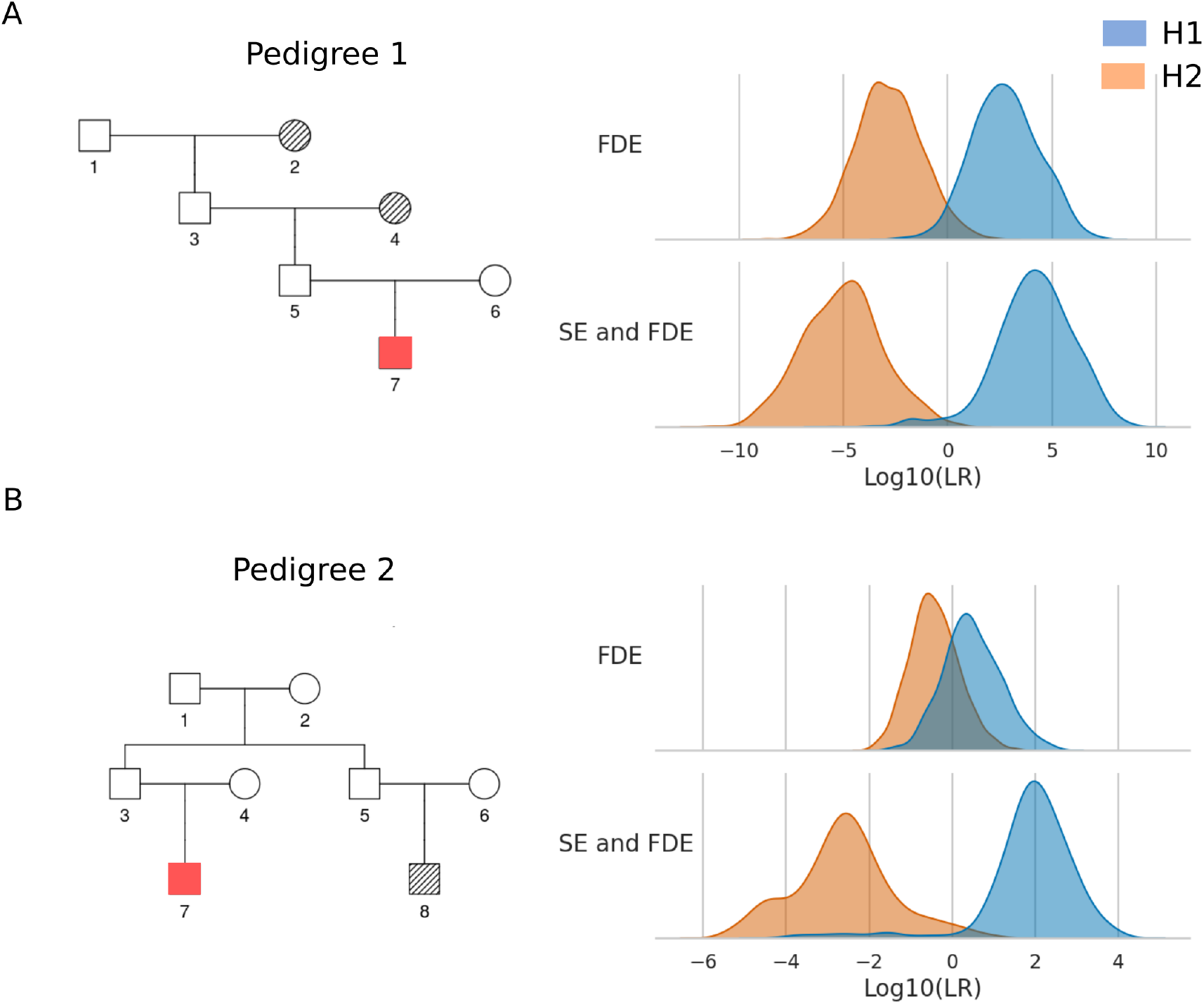
The distributions of *Log*_10_(LR) for two different pedigrees (labeled as Pedigree 1 and Pedigree 2), under two scenarios: using only the FDE (*top row*) versus combining FDE with SE (*bottom row*).

Figure 5 illustrates the distributions of *Log*_10_(LR) under two scenarios: one using FDE alone, and one where FDE is combined with SE (Section 2.4, Equation 8). In both cases, a classification threshold of LR = 1 was applied.

For Pedigree 1, the FDE only approach already showed relatively strong performance, reaching a *FNR* of 0.031 and a *FPR* of 0.039, corresponding to a *MCC* of 0.93. However, once SE was incorporated, the *FPR* dropped to 0.006, and the *MCC* rose to 0.96.

Even more importantly, improvements were observed for Pedigree 2, which initially proved difficult to resolve with FDE alone, with *FNR* = 0.246, *FPR* = 0.22 and *MCC* = 0.53. By integrating SE data, we substantially reduced both *FNR* and *FPR*, to 0.053 and 0.029, respectively, and increased the *MCC* to 0.92.

## 4 Discussion

In this work, we addressed fundamental limitations where STRs and unlinked FDE alone are not enough for reaching conclusive results: (i) cases where FDE likelihoods remain invariant under permutation, preventing discrimination of directionality; and (ii) cases where SE becomes crucial due to degraded samples or lack of close relatives. For (i), we incorporated Bayesian priors derived from empirical age distributions and intergenerational intervals (Examples 3.1), generating informative posterior probabilities for alternative pedigree configurations. For (ii), we introduced *comparison data* using transition matrices to model correspondence between phenotypic traits observed in Ups and Ms (Equations (3) and (4)).

### 4.1 Incorporation of supplementary evidence

Despite the potential value of SE in kinship analysis, formal methods for its incorporation remain largely unexplored [13]. This gap stems from (i) the sufficiency of conventional genetic evidence, (ii) preference for qualitative expert assessments over quantitative approaches, and (iii) challenges in developing robust statistical models with accurately estimated parameters for diverse supplementary evidence forms.

While argument (i) is valid for high-quality DNA contexts, missing persons investigations usually require integrative approaches considering all available evidence [6], and the inappropriate application of non-genetic filtering criteria can yield erroneous conclusions [13]. Presently, different software [22, 35] allows the filtering approach as the main procedure for SE. Moreover, multiple studies emphasise that the identification phase in missing persons investigations must adopt an integrative approach that combines different lines of evidence [12, 13, 36, 37, 38, 39].

Argument (ii) becomes problematic in large-scale databases where case-by-case expert review is logistically unfeasible. In these cases, underpowered FDE, for example, due to the lack of close relatives of the missing, can generate many false positives and negatives [1]. Usually, false positives will be checked. But, importantly, false negatives represent silent errors that preclude SE incorporation that might lead to successful resolutions [2]. Moreover, in other cases, the search can be started using only SE through large databases, where case-by-case examination also becomes difficult [8, 9, 10].

Regarding (iii), specifying SE models and parameters presents challenges compared to FDE-based approaches, which benefit from established biological foundations and extensive parameter estimation data. SE traits inferred through anthropological or DNA-based predictive models are typically accompanied by probabilistic estimates that may depend on the quality of the sample (e.g., age-at-death, height, ancestry inference) [32, 40, 41], facilitating uncertainty estimates, while traits from verbal accounts or historical documents present greater quantification challenges. However, corroborative sources such as photographs, official legal records [9, 10], and OSINT information can increase confidence [42].

In the absence of reliable error rates, we recommend sensitivity analyses by systematically varying key parameters and evaluating likelihood ratio stability across plausible ranges, paralleling FDE practices for mutational and dropout parameters [22]. We applied this approach to our SE models (Appendix C), revealing that increasing uncertainty consistently decreased LRs under *H*_1_ while increasing them under *H*_2_, confirming appropriate evidential weight adjustment, while population-rare traits generated higher LRs, reinforcing the forensic principle that trait rarity correlates with informational value [31]. Sensitivity analyses are also useful for priors settings [43].

An important limitation is that our framework models conditionally on observed missing person data, assuming certainty, yet uncertainty exists on both sides of the comparison for unidentified and missing persons alike [9, 44]. This methodological challenge remains largely unaddressed, also in traditional FDE models, which assume certainty in reference pedigree structures despite known limitations and potential errors in stated relationships between references [45]. This issue requires further methodological development and represents a promising line of research.

### 4.2 Alternative methodological approaches

Here, we examine alternative approaches and limitations that represent promising research avenues.

#### 4.2.1 Directional data

Beyond using priors to include directional SE evidence (Section 2.3), data-driven approaches are possible. Assuming age distributions for individuals A and B are normally distributed and independent, 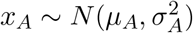 and 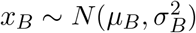, hypotheses *H*_1_ and *H*_2_ in Figure 1 are possible if *q*_1_ = *P* (*x*_*A*_ *− x*_*B*_ *> g*) and *q*_2_ = *P* (*x*_*B*_ *− x*_*A*_ *> g*), where *g* is a minimal parent-offspring age difference (e.g., *g* = 15). Using point estimates for parameters *µ*_1_, *µ*_2_, *σ*_1_, *σ*_2_, we calculate *q*_1_, *q*_2_ and *q*_3_ = 1 *− q*_1_ *− q*_2_ as priors to obtain posteriors 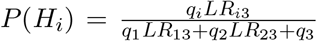, where *H*_3_ specifies A and B as unrelated. For ages estimated at 30 and 10 with standard deviations 5 and 2, respectively, and LRs *LR*_13_ = *LR*_23_ = 100000, we obtain *q*_1_ = 0.82, *q*_2_ *≈* 0, *q*_3_ = 0.18, yielding *P* (*H*_1_) *≈* 1. This integration of age-based directional evidence provides a statistical framework resolving relationship ambiguities where traditional DNA evidence alone is insufficient.

#### 4.2.2 Comparison data

The variables analysed as comparison data usually differ in modeling complexity: categorical variables (e.g., biological sex) are mathematically straightforward, while continuous variables (e.g., age) require probability distributions or fixed categories based on predetermined ranges. The floating bin approach [27] handles continuous variables by conservatively assigning matching windows accounting for measurement uncertainty. Defining likelihood models for *H*_2_ (UP is not M) requires population-based statistics assuming UP represents a randomly selected individual from the reference population. While census records provide accessible sources for age, biological sex, and similar variables, pigmentation traits present greater challenges [46], though efforts are underway to build reliable databases [34, 32].

Regarding the setting for *H*_2_, when population-specific demographic data is available, such as in closed DVI scenarios, reference frequencies for demographic variables should be adjusted to reflect the actual composition of missing and unidentified persons rather than general population frequencies, which are commonly used in open MPI scenarios [13]. However, in the absence of reliable demographic information, uncertainties should be taken into account through sensitivity analyses as previously discussed.

LRs can be combined through direct multiplication by assuming independence between characteristics, and it is reasonable since our FDE contained no phenotype-predictive markers. While independence is logical for variables such as biological sex and age, this assumption does not hold for pigmentation traits [34]. Importantly, this dependency has not been considered in previous studies [13]. Modeling dependency between variables can significantly enhance identification by capturing informative relationships that would otherwise be overlooked, with conditional dependency allowing LRs to reflect both direct trait concordance probabilities and population structure (Example 3.4). However, such dependencies are challenging to specify explicitly without exhaustive database analysis enabling reliable co-occurrence computation (non-parametric approach), though parametric approaches are possible with smaller databases (Appendix A).

Finally, we anticipate that broader adoption of massively parallel sequencing (MPS) technologies will increase the availability of DNA-based phenotyping and enhance statistical power in kinship inference by accessing denser SNP data [18, 41]. Nonetheless, distant-relative inference will continue to involve substantial uncertainty even with these approaches [47]. Consequently, the expansion of MPS will generate new methodological and statistical challenges regarding the incorporation and formalization of supplementary evidence in genealogical and forensic research.

## 5 Conclusion

Our framework integrates forensic DNA evidence with supplementary evidence through a statistical approach that addresses kinship analysis limitations. We formalize supplementary evidence using Bayesian priors for directional data and transition matrices for comparison data. The open-source `mispitools` package enables forensic practitioners to apply this framework. This work demonstrates that properly modeled supplementary evidence provides discriminatory information for kinship analyses, offering solutions for challenging scenarios from standard paternity testing to large-scale disaster victim identification where resolutions might otherwise prove elusive.

## Compliance with Ethical Standards

Not applicable.

## Research involving human participants, their data or biological material

Not applicable.

## Informed consent

Not applicable.

## Data Availability Statement

For access to BALSAC genealogical data used in the simulations, please contact: https://balsac.uqac.ca/en/contact/. The open source code for the analyses is available at: https://github.com/MarsicoFL/mispitools.

## Authors’ Contributions

TE: Conceptualization, Methodology, Software, Formal analysis, Writing - Original Draft, Writing - Review & Editing. FM: Conceptualization, Methodology, Software, Formal analysis, Writing - Original Draft, Writing - Review & Editing

## A Parametric dependence model

In general, there appears not to be any general parametric models for dependence suitable for our applications. However, in the simple case with two binary variables a model can be established and studied. Let *P* (*x*_1_ = 1) = 1 *− P* (*x*_1_ = 0) = *p* and *P* (*x*_2_ = 1) = 1 *− P* (*x*_2_ = 0) = *q* and consider the joint distribution given in Table 4. The main advantages of this parametric distribution are that we can estimate *θ* from data and study the impact of dependence as a function of *θ*. Note that

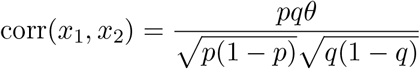

**Table 4:** Joint bivariate distribution. Note that admissible values for *θ* are those leading to a proper probability distribution.

| $x_1/x_2$ | 0 | 1 | sum |
| --- | --- | --- | --- |
| 0 | $1 - p - q + pq(1 + \theta)$ | $q(1 - p(1 + \theta))$ | $1 - p$ |
| 1 | $p(1 - q(1 + \theta))$ | $pq(1 + \theta)$ | $p$ |
| sum | $1 - q$ | $q$ | $1$ |

If *θ >* 0 there is positive dependence and then *L*_*dep*_(0, 0) *> L*_*ind*_(0, 0) and *L*_*dep*_(1, 1) *> L*_*ind*_(1, 1) whereas the opposite inequalities apply for (1,0) and (0,1). This allows us to conclude on the impact of dependence depending on whether *θ >* 0 or not. For instance, the ratio of LR assuming independence to the one for dependence is (1 + *θ*) when *x*_1_ = *x*_2_ = 1.

## B Balsac genealogies

Our analysis is based on a cohort of 2,077 individuals, with a nearly balanced sex distribution (50.5% male [n = 1,049] and 49.3% female [n = 1,023]). Analysis of mortality patterns yields a mean age at death of 47.2 years and a median of 55 years, with 25% of individuals succumbing before the age of 18. Despite evidence of elevated early-life mortality, the majority of deaths occur between 55 and 72 years, and the maximum observed lifespan reaches 102 years (Figure S1A). The birth years ranged from 1665 to 1822 (Figure S1B). Examination of half-sibling relationships reveals a marked asymmetry: 62.5% (n = 346) of half-sibling pairs share a paternal lineage, while only 37. 5% (n = 208) are maternal half-siblings. For anonymity purposes, in the implemented models in mispitools we do not add the real datasets, but only summary metrics (frequency distribution of traits) obtained from it.

**Figure S1:**
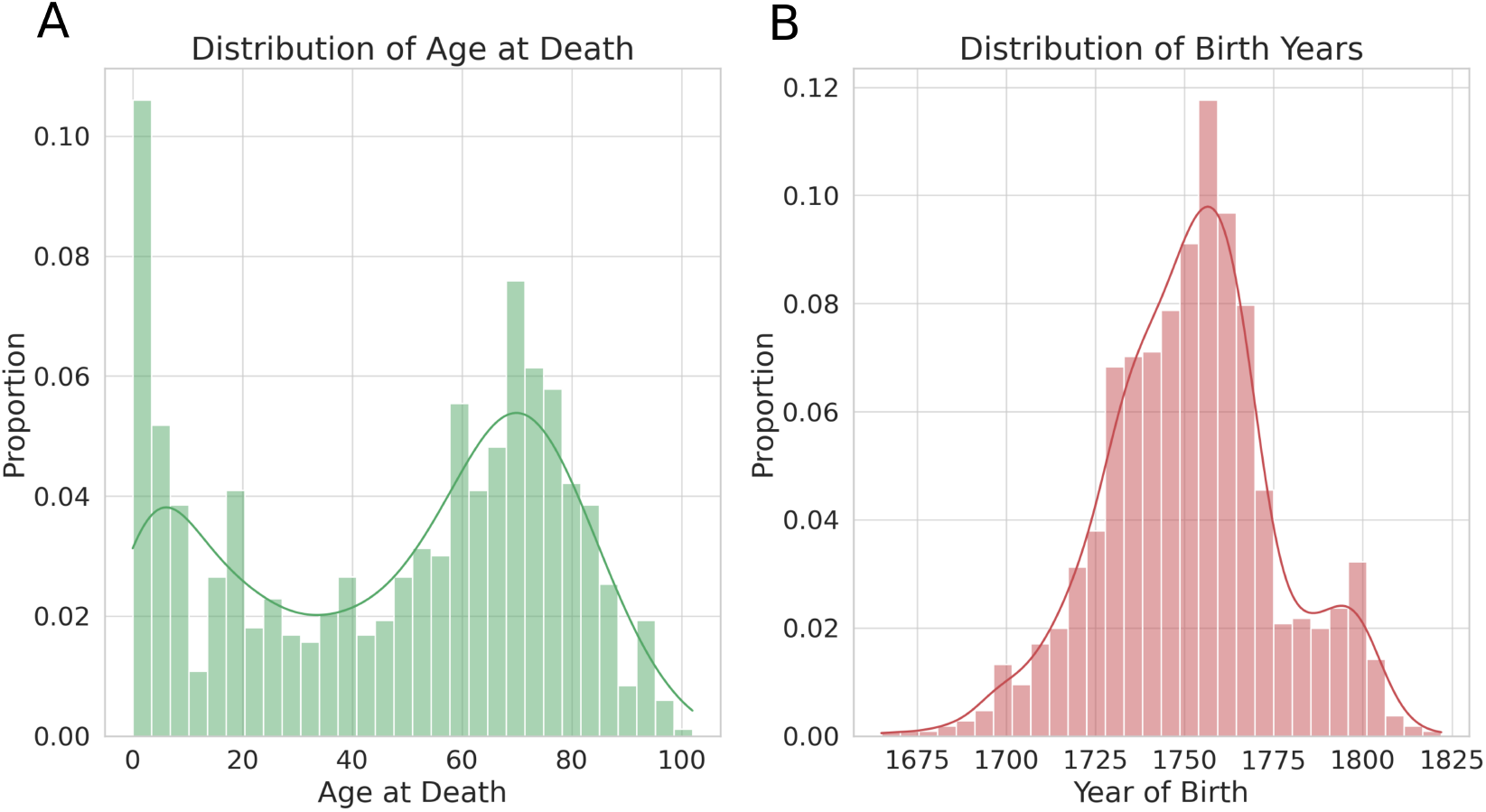
Demographic and genealogical distributions from the BALSAC database. Panel (A) Distribution of Age at death, (B) Year of birth distribution.

## C Sensitivity analysis

To evaluate the robustness of the LR models, we performed a sensitivity analysis systematically varying key model parameters. In the sex-based analysis (Figure S2A), the error parameter (*ε*) was varied from 0.01 to 0.4 while the female population proportion was adjusted from 0.1 to 0.4. For the LR based on age (Figure S2B), with the missing person age (MPa) fixed at 40 and the simulation parameters kept constant (that is, *γ* = 0.07 and *ε*_age_ = 0.05), the age error range, or also called the matching window, varied from 1 to 20. In the hair colour analysis (Figure S2C), the baseline hair colour error (*ε*) varied from 0.01 to 0.4 and the population proportion for hair colour 1 (p1) varied between 0.1 and 0.2 (with the remaining proportions scaled proportionally to sum to 1). In each scenario, 100,000 simulations were performed under both hypotheses: *H*_1_ (the unidentified individual is the missing person) and *H*_2_ (the unidentified individual is not the missing person), and the median LR was calculated.

The results reveal a clear trend: as the uncertainty increases (for example, with higher error rates), the LR values under *H*_1_ decrease while those under *H*_2_ increase. This behavior indicates that the model effectively captures the diminishing weight of the evidence as the uncertainty grows. In addition, a similar pattern is observed when the specific trait value (female in Figure S2A and the selected hair colour in Figure S2C) becomes more prevalent. This finding reinforces a well-known principle in forensic analysis: When the observed trait is less common, it is more informative and contributes greater weight to the evidence.

**Figure S2:**
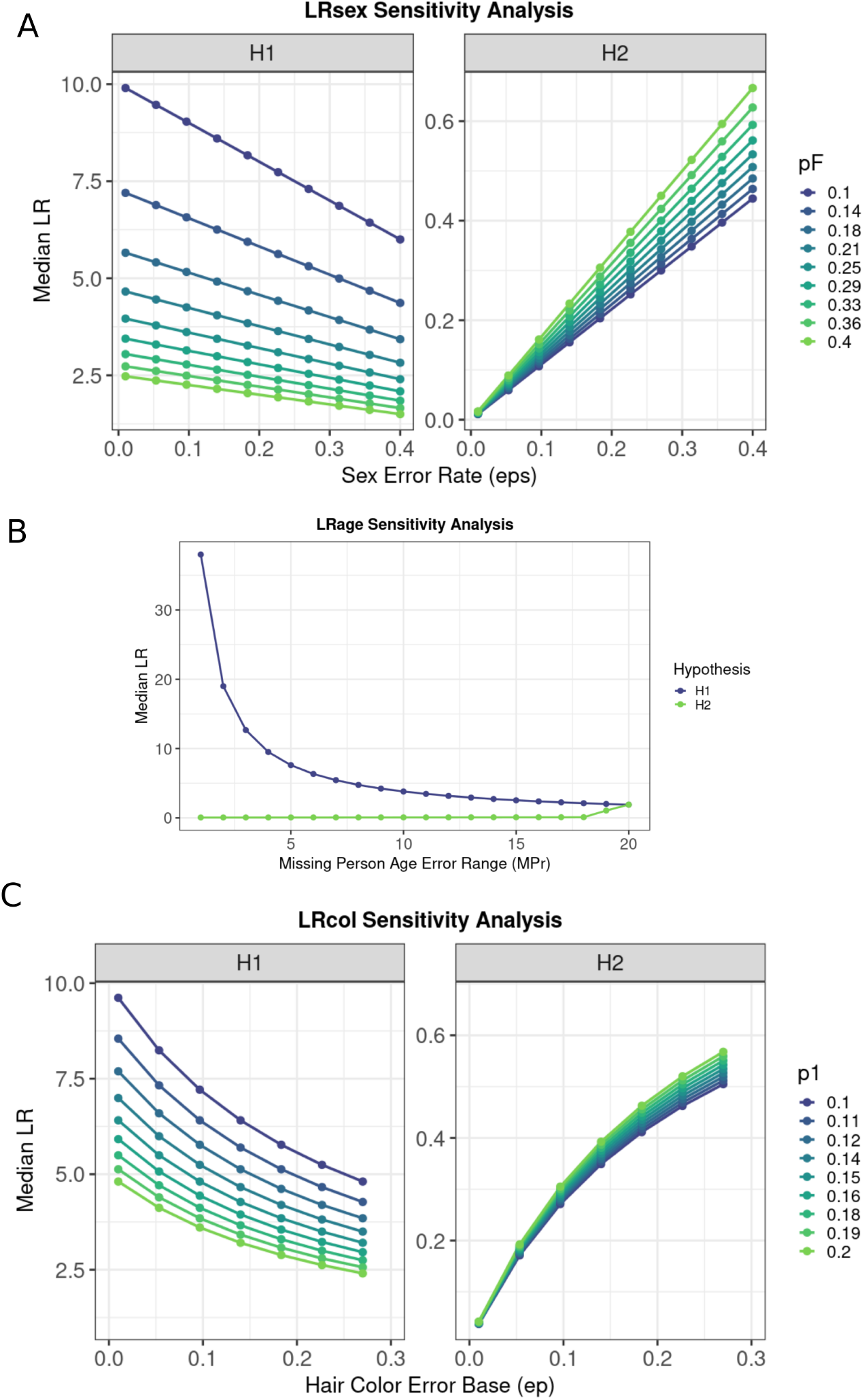
Sensitivity analysis of likelihood ratio estimates for (a) sex, (b) age, and (c) hair colour. For LRsex, the error rate (*ε*) and female population proportion (pF) were varied from 0.01–0.4 and 0.1–0.4, respectively. For LRage, with age fixed at 40, the age error range was varied from 1 to 20. For LRcol, the hair colour error (*ε*) ranged from 0.01 to 0.4 and the proportion for hair colour 1 (p1) from 0.1 to 0.2. Median LR values (computed from 500 simulations) are displayed for both hypotheses (*H*_1_ and *H*_2_), illustrating the impact of parameter uncertainty on LR estimates.

