## supplementary code for "Using all available evidence to solve kinship cases": usingAll.R

```
# Setting up

install.packages("mispitools")
install.packages("pedtools")
library(mispitools)
library(pedtools)
library(forrel)

# Section 1: SE data
## Analyzing comparison data based LRs

### Section 1.1: Biological sex
sex_H1 <- LRsex("F", H = 1, LR = TRUE, seed = 123, nsims = 500)
sex_H2 <- LRsex("F", H = 2, LR = TRUE, seed = 123, nsims = 500)
sex_LRs <- as.data.frame(cbind(sex_H2$LRs, sex_H1$LRs))
names(sex_LRs) <- c("Unrelated", "Related")
sex_LRs

LRdist(sex_LRs)
Trates(sex_LRs, 1)

### Section 1.2: Age
age_H1 <- LRage(40, H = 1, LR = TRUE, seed = 123, nsims = 500)
age_H2 <- LRage(40, H = 2, LR = TRUE, seed = 123, nsims = 500)
age_LRs <- as.data.frame(cbind(age_H2$LRa, age_H1$LRa))
names(age_LRs) <- c("Unrelated", "Related")
age_LRs

LRdist(age_LRs)
Trates(age_LRs, 1)

### Section 1.3: Pigmentation trait
col_H1 <- LRcol(1, H = 1, LR = TRUE, seed = 123, nsims = 500)
col_H2 <- LRcol(1, H = 2, LR = TRUE, seed = 123, nsims = 500)
col_LRs <- as.data.frame(cbind(col_H2$LRc, col_H1$LRc))
names(col_LRs) <- c("Unrelated", "Related")
col_LRs

LRdist(col_LRs)
Trates(col_LRs, 1)

### Section 1.4: Combined comparison data
comb_H1 <- sex_H1$LRs * age_H1$LRa * col_H1$LRc
names(comb_H1) <- "LRH1"
comb_H2 <- sex_H2$LRs * age_H2$LRa * col_H2$LRc
names(comb_H2) <- "LRH2"
comb_LRs <- as.data.frame(cbind(comb_H2, comb_H1))
names(comb_LRs) <- c("Unrelated", "Related")

LRdist(comb_LRs)
Trates(comb_LRs, 1)

### Section 1.5: Visualization, exploring overall likelihoods and LRs
POPl <- CPT_POP(propS = c(0.5, 0.5),
                MPa = 40,
                MPr = 6,
                propC = c(0.3, 0.2, 0.25, 0.15, 0.1))

### Note: Population frequencies are obtained from the reference datasets

MPl <- CPT_MP(MPs = "F", MPc = 1,
              eps = 0.05, epa = 0.05,
              epc = Cmodel())

MPl/POPl

CondPlot(MPl,POPl)


# Section 2: FDE data
## Simulating DNA-based LRs in underpowered cases

set.seed(1234)
f <- getfreqs(Argentina)
ped1 <- linearPed(3)
ped1 <- setMarkers(ped1,locusAttributes = f)
ped1 <- profileSim(ped1,N = 1,ids = c(2,4))
datasimx = simLRgen(ped1, missing = 7, numsims = 500, seed = 1234)
plot(ped1, hatched = typedMembers(ped1), cex = 0.9)

LRdist(datasimx)
Trates(datasimx, threshold = 1)

ped2 <- cousinPed(1)
ped2 <- setMarkers(ped2,locusAttributes = f)
ped2 <- profileSim(ped2,N = 1,ids = 8)
datasimy = simLRgen(ped2, missing = 7, numsims = 500, seed = 1234)

plot(ped2, hatched = typedMembers(ped2), cex = 1)


LRdist(datasimy)
Trates(datasimy, threshold = 1)


# Section 3: Combining FDE and SE data
## Combining DNA-based LRs and comparison data based LRs
#### Note: We consider the same SE data for both FDE cases

datasimx2 <- simLR2dataframe(datasimx)
combinedx_H1 <- datasimx2$Related * sex_H1$LRs * col_H1$LRc * age_H1$LRa
combinedx_H2 <- datasimx2$Unrelated * sex_H2$LRs * col_H2$LRc * age_H2$LRa
combined_datasimx <- as.data.frame(cbind(combinedx_H2, combinedx_H1))
names(combined_datasimx) <- c("Unrelated", "Related")
combined_datasimx

LRdist(combined_datasimx)
Trates(combined_datasimx, 1)


datasimy2 <- simLR2dataframe(datasimy)
combinedy_H1 <- datasimy2$Related * sex_H1$LRs * col_H1$LRc * age_H1$LRa
combinedy_H2 <- datasimy2$Unrelated * sex_H2$LRs * col_H2$LRc * age_H2$LRa
combined_datasimy <- as.data.frame(cbind(combinedy_H2, combinedy_H1))
names(combined_datasimy) <- c("Unrelated", "Related")
combined_datasimy

LRdist(combined_datasimy)
Trates(combined_datasimy, 1)
```
